# Neuroglian regulates *Drosophila* intestinal stem cell proliferation through enhanced signaling via the Epidermal Growth Factor Receptor

**DOI:** 10.1101/2020.11.17.385781

**Authors:** Martin Resnik-Docampo, Kathleen M. Cunningham, S. Mateo Ruvalcaba, Charles Choi, Vivien Sauer, D. Leanne Jones

## Abstract

The *Drosophila melanogaster* intestine is an excellent system for elucidating mechanisms regulating stem cell behavior under homeostatic conditions or in response to injury, stress, or ageing. Here we show that the septate junction (SJ) protein Neuroglian (Nrg) is expressed in intestinal stem cells (ISCs) and daughter enteroblasts (EBs) within the fly midgut, the equivalent of the mammalian small intestine. Although Nrg localizes to the plasma membrane, SJs are not present between ISC/EBs, suggesting Nrg plays a different role in this tissue. Generation of ISCs homozygous for a null allele of *Nrg* revealed that Nrg is required for ISC proliferation in young flies, and depletion of Nrg from ISCs/EBs was able to suppress the increase in ISC proliferation with age. Conversely, overexpression of *Nrg* in ISC/EBs was sufficient to drive ISC proliferation, leading to an increase in cells expressing ISC/EB markers. In addition, we observed an increase in EGFR activation. Genetic epistasis experiments revealed that Nrg acts upstream of EGFR in the midgut to regulate ISC proliferation. As Nrg function is highly conserved in mammalian systems, our work characterizing the role of Nrg in the intestine has implications for the etiology and treatment of intestinal disorders due to altered ISC behavior.

## Introduction

Adult stem cells (SCs) maintain tissue homeostasis through the balanced generation of new daughter stem cells and progenitor cells destined to differentiate. In addition, adult stem cells serve as a reservoir of cells for repair of tissues and organs after damage. Studies have shown that age-related changes in SC function likely lead to loss of homeostasis over time and may contribute to age-onset disease (Jones and Rando, 2011). Therefore, understanding the mechanisms involved in regulating stem cell behavior and how these mechanisms are altered with age will uncover therapeutic targets for regenerative medicine in order to treat age-onset and/or degenerative diseases.

The *Drosophila* midgut, the functional equivalent to the mammalian small intestine, is maintained over time by resident intestinal stem cells (ISCs) (Micchelli and Perrimon, 2006; Ohlstein and Spradling, 2006). The ISCs are multipotent and can divide to produce more ISCs or enteroblasts (EBs) that differentiate into absorptive enterocytes (ECs) or secretory enteroendocrine cells (EEs), all of which are needed to maintain homeostasis. Additional reports suggest that ISCs can differentiate into EE cells directly, without progressing through the EB state (Amcheslavsky et al., 2014; Biteau and Jasper, 2014; Guo and Ohlstein, 2015; Zeng and Hou, 2015). Several highly conserved signaling pathways, including the Epidermal Growth Factor Receptor (EGFR) and Notch (N) pathways, play essential roles in regulating ISC proliferation and differentiation, respectively (Li and Jasper, 2016; Nászai et al., 2015). The Notch pathway, in particular, is crucial for regulating the balance of differentiated EC and EE cells: high N signaling in ISCs leads to an EC fate, while low/no N signaling leads to production of EE cells (Bardin et al., 2010; Ohlstein and Spradling, 2007; Perdigoto et al., 2011; Reiff et al., 2019). EGFR/Ras/MAPK signaling acts as a permissive signal for proliferation in ISCs, coordinating with JAK/STAT signaling to regulate ISC growth and division (Biteau and Jasper, 2011; Buchon et al., 2010; Cordero et al., 2012; Jiang et al., 2009, 2011; Jin et al., 2015; Xu et al., 2011). EGF ligands and the EGFR itself are responsive to additional, conserved homeostatic or stress signals integrated from the environment (Buchon et al., 2010; Cordero et al., 2012; Du et al., 2020; Ngo et al., 2020; Zhang et al., 2019). Remarkably, similar observations have been made in the mammalian intestine, underscoring the usefulness of the fly intestine as a model to uncover conserved mechanisms regulating ISC behavior (Jasper, 2020).

A number of age-related changes occur in the *Drosophila* intestine, such as increased ISC proliferation, accumulation of EB-like cells that express stem cell markers as well as hallmarks of differentiated cells, bacterial dysbiosis, and loss of the intestinal barrier (Jasper, 2020). In flies and mammals, disruption of intestinal barrier function and increased intestinal permeability correlate with compromised integrity of cell-cell junctions, known as occluding junctions (Marchiando et al., 2010; Rera et al., 2012; Resnik-Docampo et al., 2017; Vancamelbeke and Vermeire, 2017). These specialized structures-tight junctions (TJs) in vertebrates and septate junctions (SJs) in arthropods- are important for regulating paracellular flow between apical and basal epithelial surfaces. In a previous study investigating age-related changes to the intestinal barrier, we found that the SJ protein Neuroglian (Nrg) is strongly expressed in the hindgut (Resnik-Docampo et al., 2017). However, we also observed expression within ISCs/EB ‘nests’ in the midgut. Here, we describe a role for Nrg in regulating ISC behavior through potentiation of signaling via EGFR in both young and aged flies.

## Results

### Nrg is expressed in ISC/EB nests in the *Drosophila* posterior midgut

Our lab previously reported expression of known SJ proteins in the *Drosophila* intestine and described how expression and localization patterns change as a consequence of aging (Resnik-Docampo et al., 2017). While the SJ protein Nrg was strongly expressed in the pleated septate junctions in the *Drosophila* hindgut (Fig. 1A, B-B”), Nrg protein was also detected in ISC/EB ‘nests’ that express the canonical ISC/EB marker Esg (Fig. 1A, C-C”, Fig S1A). In contrast, no Nrg protein was detected in ECs within the midgut, consistent with previous observations (Baumann, 2001).

**Figure 1.**
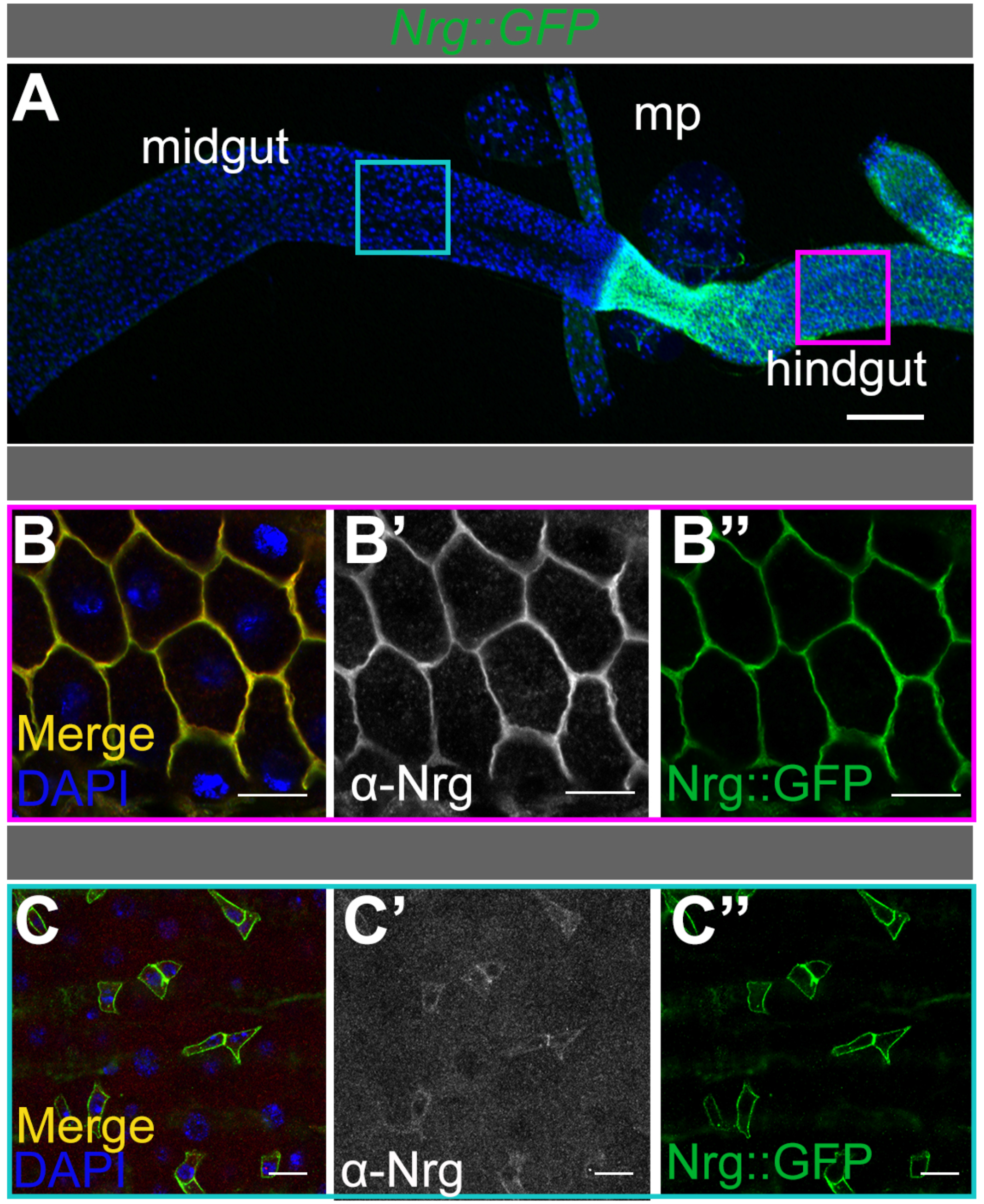
Neuroglian (Nrg) is expressed in the ISCs/EBs of the *Drosophila* midgut. A) Representative low magnification image of Nrg::GFP (green) expression in the adult gut including the midgut, malpigian tubules (mp), and hindgut. Scale bar = 100μm B and C) Adult hindgut (B, magenta border) and midgut (C, teal border) of an adult expressing *Nrg::GFP* (B, B” and C, C”) and stained with anti-Nrg antibody (B, B’ and C, C’) (See Methods). Note expression of Nrg in ISC/EB “nests” of the midgut. Scale bar = 20μm.

Two different protein isoforms of Nrg exist that differ at the C-terminal cytoplasmic domain: a neuronal-specific isoform, Nrg^180^, and another isoform, Nrg^167^, that is more generally expressed (Hortsch et al., 1990). In order to confirm that a *Nrg::GFP* fusion protein accurately reflects Nrg protein expression, we generated an antibody that detects both Nrg isoforms (see Methods); anti-Nrg antibody specificity was confirmed using depletion of Nrg via RNAi in wing imaginal discs (Supplementary Fig 1B-C’). Using the antibody, confocal immunofluorescence (IF) microscopy confirmed endogenous Nrg expression and localization patterns in the hindgut and midgut (Fig. 1A, B-B’, C-C’). Consistent with our observations, recent single cell sequencing data profiling of the *Drosophila* midgut found *Nrg* among the most enriched genes in ISC/EB clusters (Hung et al., 2020). In addition, RNA-seq analysis performed from 5-day-old flies (Resnik-Docampo et al., 2017) revealed that the *Nrg^167^* isoform is the primary isoform expressed in the posterior midgut, while the neuronal *Nrg^180^* isoform was detected at very low levels (Supplementary Fig. 1D).

As SJs are restricted to EC-EC and EC-EE junctions in the midgut (Resnik-Docampo et al., 2017), these data suggested that Nrg is likely not acting as a SJ protein in the *Drosophila* midgut. In addition to its role as a SJ protein, Nrg is one example of the cell adhesion molecules (CAMs) that play a role in the developing nervous system (Enneking et al., 2013; Goossens et al., 2011; Kristiansen et al., 2005; Kudumala et al., 2013; Moscoso and Sanes, 1995). In this context, Nrg has been demonstrated to modulate EGFR/FGFR signaling in order to regulate axon extension and guidance in sensory neurons (García-Alonso et al., 2000; Islam et al., 2003; Nagaraj et al., 2009). In mammals, the role of the Nrg homolog, L1CAM, is conserved in nervous system development (Dahme et al., 1997; Godenschwege et al., 2006; Jouet et al., 1994; Kudumala et al., 2013; Schäfer and Altevogt, 2010), and L1CAM interactions with EGFR/FGFR are also preserved in mammalian cells (Donier et al., 2012; Islam et al., 2004; Kulahin et al., 2008). Interestingly, human L1CAM (hL1CAM) expression rescues *Nrg* loss-of-function phenotypes in *Drosophila*, demonstrating a remarkable conservation of function (Godenschwege et al., 2006; Kristiansen et al., 2005; Kudumala et al., 2013).

### *Nrg* is required for ISC proliferation in the posterior midgut

In order to investigate the role of Nrg in the posterior midgut, FRT-mediated recombination was used to generate positively marked (GFP^+^) ISC ‘clones’ homozygous mutant for either a null allele of *Nrg*, *Nrg^14^* (Enneking et al., 2013) or a strong hypomorphic allele, *Nrg^G00413^* (Fig. 2A,C). Wild type, GFP^+^ control clones were generated in parallel. Quantification of the number of clones per gut 7 days post-clone induction revealed no difference in frequency when comparing wild type with *Nrg^14^* or *Nrg* mutant clones, suggesting that *Nrg* mutant ISCs are not lost (Fig. 2B). However, detailed analysis of the clonal cell population showed that *Nrg^14^* or *Nrg*^G00413^ clones often consisted of single ISCs or EBs (91.0% and 94.2%, respectively), when compared to controls (63.7%) (Fig 2A, CD). The increase in single cell clones corresponded to a reduction in clones containing differentiated cells, with 8.2% of *Nrg^14^* clones containing only EEs or ECs, compared to 24.4% for controls. Furthermore, the number of clones containing all the cell types dropped from 10.3% in controls to 0.8% in *Nrg^14^* mutants and 0% in *Nrg^G00413^* mutants (Fig 2D). Generation of ISC/EB clones expressing an RNAi construct targeting *Nrg* resulted in a similar shift to clones containing single cells (Supplementary Figure 2B-B’, F, G) after 14 days, when compared to controls (Supplementary Figure 2A-A’, D-F, G). However, there was no decrease in the number of GFP+ clones after 7 or 14 days (Supplementary Figure 2C, E). Taken together, these data suggest that Nrg plays a role in regulating ISC proliferation.

**Figure 2.**
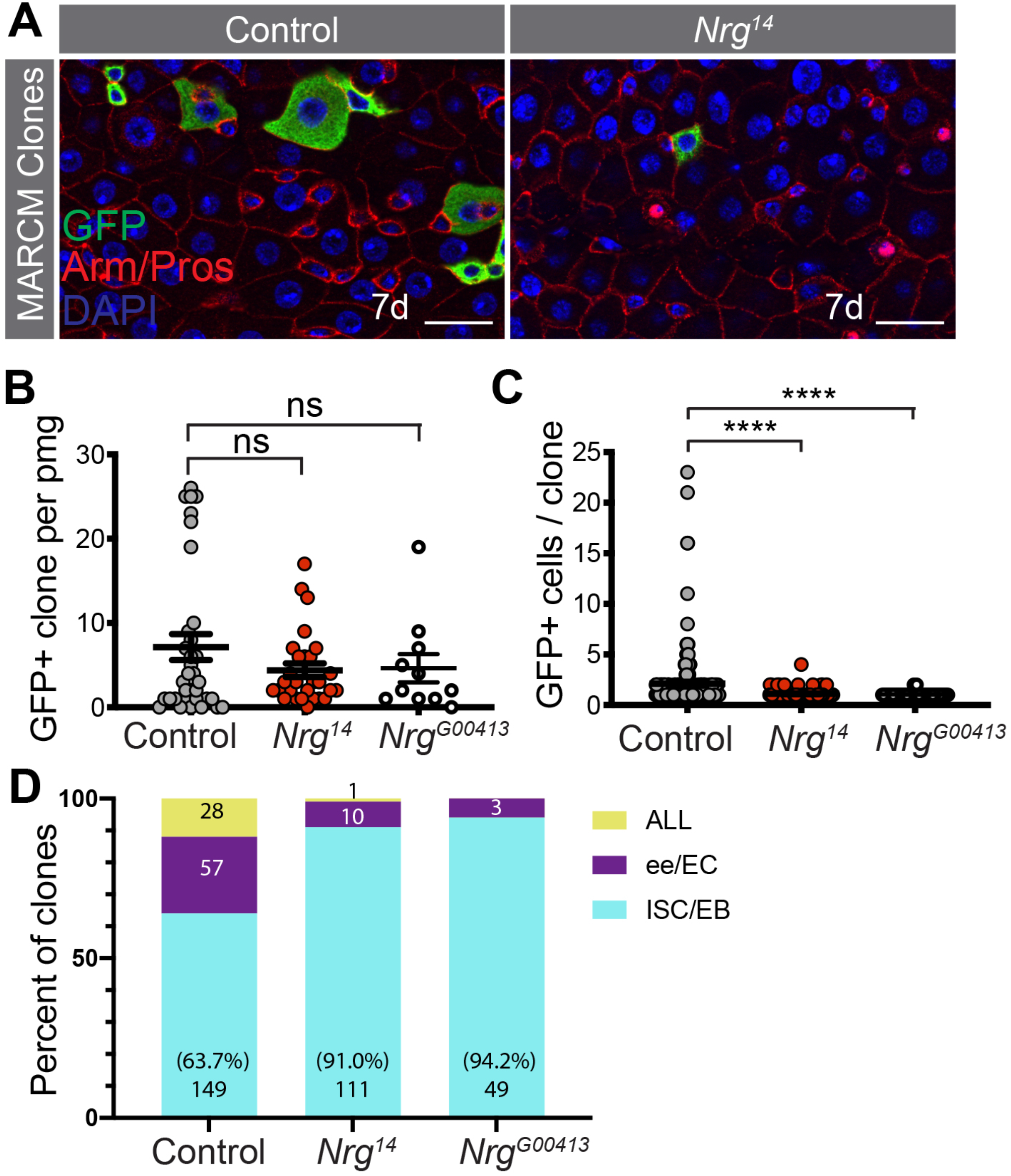
Nrg is required for ISC proliferation. A) Examples of midguts 7 days after FRT-mediated clonal generation in control *(hsflp FRT19A, tubGal80^ts^ / FRT19A; UAS-GFP/ tubGa/4)* and *Nrg^14^ (hsflp FRT19A, tubGal80^ts^ / FRT19A Nrg^14^; UAS-GFP/ tubGa/4*backgrounds. Clones are positively marked in green (See methods). B) Total number of GFP+ cells (MARCM clones) per posterior midgut in control, *Nrg^14^*, and *Nrg^G00413^* backgrounds, N= 34 control, 28 *Nrg^14^*, 11 *Nrg^G0041^* guts. C) Quantification of number of GFP+ cells per clone in 7 days after FRT-mediated clonal generation in control, *Nrg^14^*, and *Nrg^G00413^* backgrounds. N = 225 control, 122 *Nrg^14^*, 52 *Nrg^G0041^* clones. D) Characterization of the type of GFP+ cells (ISCs/EBs, enteroendrocine cells(EEs)/enterocytes (ECs), or all types) in A N = 225 control, 122 *Nrg^14^*, 52 *Nrg^G0041^* clones. Scale bar = 20μm.

### *Nrg* over-expression in ISC/EBs induces ISC proliferation

As Nrg appeared to be required for ISC proliferation, we wanted to determine whether targeted over-expression of *Nrg* in ISCs/EBs was sufficient to induce proliferation. To do so, a construct encoding *Nrg^167^* was overexpressed utilizing an RU486-inducible Gene-Switch ‘driver’ line that is expressed in ISCs and EBs *5961^GS^* (see Materials and Methods). Addition of RU486 to food (RU+) leads to induction of transgene expression, while lack of RU486 (RU-) and outcrossed controls lacking the transgene served as negative controls. Mitotic cells were detected and quantified by staining for phosphorylated histone H3 (pH3); as ISCs are the only dividing cells in the intestine, quantification of proliferation serves as a surrogate marker for the presence and activity of ISCs. After 7 days of exposure to RU486, a statistically significant increase in the number of mitotic cells was observed upon *Nrg^167^* expression, when compared to controls (Fig. 3A-B). Accordingly, we observed an increase in the number of cells expressing the ISC/EB marker Esg (Fig. 3A, C). In previous studies in the *Drosophila* nervous system, the human homolog of Nrg, hL1CAM, was found to rescue neurodevelopmental defects in *Nrg* mutants (Godenschwege et al., 2006; Kakad et al., 2018; Kudumala et al., 2013). Therefore, we wanted to test whether human hL1CAM expression was also sufficient to induce ISC proliferation. Indeed, ectopic expression of hL1CAM in ISC/EBs with the*5961^GS^* ‘driver’ also showed a significant increase in ISC proliferation, similar to *Nrg^167^* (Fig. 3A-C).

**Figure 3.**
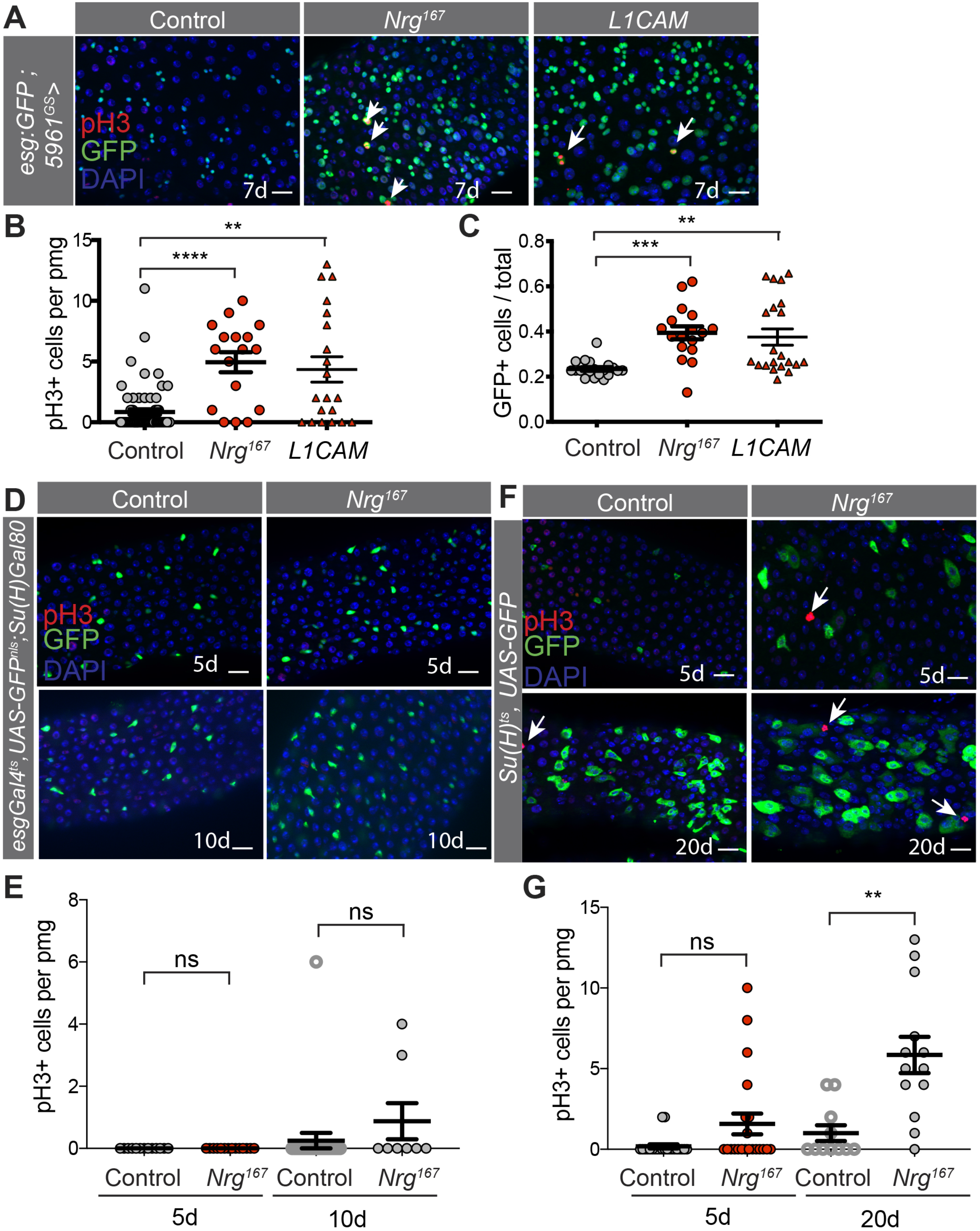
Over-expression of Nrg in ISCs/EBs induces ISC proliferation. A) Representative images from adult midguts of Control *(OreR), Nrg^167^*, and *hL1CAM* driven by *5961^GS^* for 7 days. ISC/EBs (esg:*GFP*, green), mitotic cells (pH3, red), and nuclei (DAPI, blue) are shown. Scale bar = 20μm. B) Quantification of number of mitotic cells per posterior midgut marked by pH3 per two fields of view in the pmg in A. N = 20 control, 17 *Nrg^167^*, 21 *hL1CAM* guts. C) Quantification of the number of *esg*:GFP+ cells per pmg in A. D) Representative images from adult midguts of Control *(OreR)* and *Nrg^167^* driven in ISCs only *(esgGal4, UAS-2xYFP; Su(H)Gal80, tubGal80^ts^)* for 5 or 10 days. ISCs (*GFP*, green), mitotic cells (pH3, red), and nuclei (DAPI, blue) are shown. E) Quantification of number of mitotic cells per posterior midgut marked by pH3 in C. N= 13 control 5do, 14 *Nrg^167^* 5do, 24 control 10do, 8 *Nrg^167^* 5do guts, Kruskal-Wallis test with Dunn’s multiple comparisons. F) Representative images from adult midguts of Control *(OreR)* and *Nrg^167^* driven in EBs only *(Su(H)Gal4, UAS-GFP; tubGal8C^ts^)* for 5 or 20 days. EBs *(GFP*, green), mitotic cells (pH3, red), and nuclei (DAPI, blue) are shown. G) Quantification of number of mitotic cells per posterior midgut marked by pH3 in E. N = 22 control 5do, 21 *Nrg^167^* 5do, 11 control 20do, 13 *Nrg^167^* 20do, Kruskal-Wallis test with Dunn’s multiple comparisons. Scale bar = 20μm.

Next, we wanted to distinguish whether Nrg expression in ISCs *or* EBs was sufficient to drive ISC proliferation. ISC-specific overexpression of Nrg for up to 10 days did not lead to an increase in ISC proliferation (Fig. 3D-E), as measured by pH3 staining. By contrast, Nrg overexpression in EBs only, using the *Su(H)Gal4, UAS-GFP; tubGal80^ts^*[referred to as *Su(H)^ts^]*, strongly induced ISC proliferation (Fig 3F-G). In addition, we observed an increase in EB-like, *Su(H)Gal4>GFP+* cells, similar to what is observed when Nrg is overexpressed in ISCs and EBs simultaneously (Fig. 3F). Taken together, these data indicate that Nrg expression in EBs can act in a non-autonomous manner to promote ISC proliferation.

### Mis-expression of *Nrg* contributes to age-related changes in the intestine

Aging results in an increase in ISC proliferation and an accumulation of EB-like cells that express hallmarks of both ISC/EBs and differentiating ECs (Biteau et al., 2008; Jiang et al., 2009; Li and Jasper, 2016; Park et al., 2009). Due to the increases in ISC proliferation and *esg*-expressing cells resulting from *Nrg* overexpression, we hypothesized that Nrg accumulation might contribute to age-related changes in the midgut. Analysis of RNA-sequencing data of old vs. young fly midguts (Resnik-Docampo et al., 2017) revealed that *Nrg* expression was 1.834+ 0.05 fold (p = 1.61*10^-27^) higher in midguts dissected from 45do flies than in the midguts of young flies. Consistent with the observed localization in ISC/EB nests and an expansion of EB-like cells with age, the number of cells expressing Nrg was also increased in intestines from aged flies (Fig 4B-B’, C), when compared to young controls (Fig 4A-A’, C).

**Figure 4.**
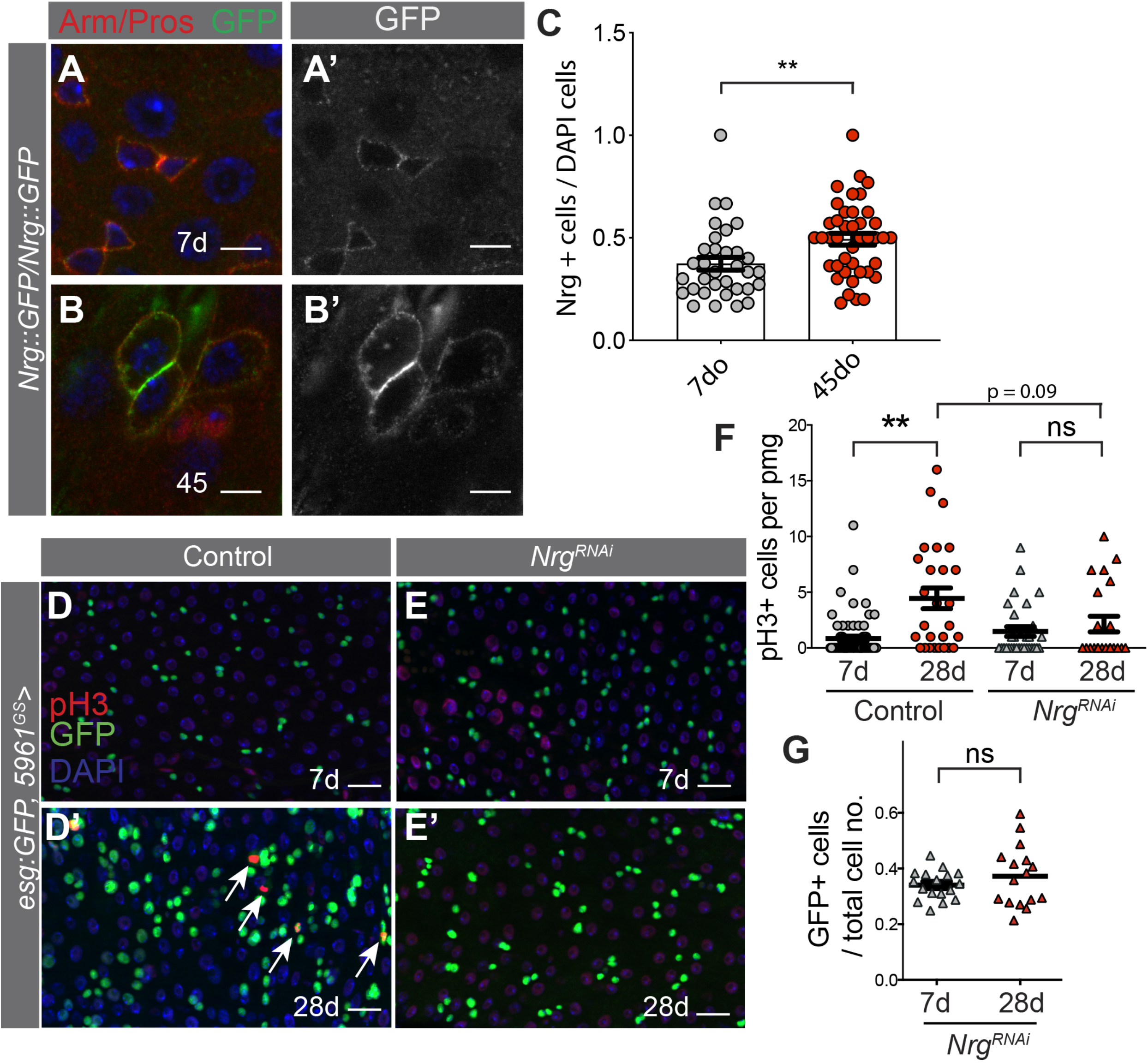
Mis-expression of Nrg contribute to age-related changes in the intestine. Representative images of Nrg::GFP in young (7d, A, A’) and old (45d, B, B’) midguts. Scale bar = 5 μm. C) Quantification of the number of Nrg+ cells per total cell number in A, N= 33 young, 41 aged fields of view (fov), unpaired two-tailed t-test. D-E) Representative images from adult midguts of Control (*OreR*, D-D’) or *Nrg^RNAl V20^* (E-E’) driven by *5961^GS^* for 7 days (young) or 28 days (old). ISC/EBs *(esg:GFP*, green), mitotic cells (pH3, red), and nuclei (DAPI, blue) are shown. F) Quantification of number of mitotic cells marked by pH3 in A per pmg N= 71 control 7do, 27 control 28 do, 30 *Nrg^RNAi^* 7do, 22 *Nrg^RNAi^* 7do guts, Kruskal-Wallis test with multiple comparisons. G) Quantification of the total number of ISCs/EBs *(esg:GFP*, green) in *5961^GS^>Nrg^RNAi^* animals shown in E-E’ N= 30 *Nrg^RNAi^* 7do, 22 *Nrg^RNAi^* 28do guts, unpaired t-test, two-sided.

Based on the observation that Nrg is required for ISC proliferation in young flies (Fig 2, Fig S2), we hypothesized that reducing *Nrg* expression in ISCs/EBs would suppress age-related increases in ISC proliferation. An increase in the number pH3 positive cells was observed in midguts dissected from 28do flies, when compared to midguts from young 7do controls, as expected (Fig. 4D-D’, F). Targeted depletion of *Nrg* expression in ISC/EBs by *5961^GS^* for 28 days blocked the age-associated increase in proliferation (Fig. 4E-E’, F). Importantly, there was no reduction in total *esg*:GFP+ cells over time when *Nrg^RNAi^* was expressed, consistent with our clonal analysis data indicating Nrg is not required for ISC maintenance and that the lack of an increase in ISC proliferation is not due to a reduction in the number of ISC/EBs (Fig 1A-B, Fig. 4G). These data demonstrate that depletion of *Nrg* from ISC/EBs is sufficient to suppress age-related increases in ISC proliferation and the accumulation of EB-like cells.

Given that *Nrg* overexpression in EBs was sufficient to drive ISC proliferation, we hypothesized that endogenous *Nrg* expression in the EB-like cells that accumulate with age would be important in driving age-related phenotypes in the gut. Therefore, we depleted *Nrg* in EBs for 20 days using *Su(H)^ts^* driver line. Indeed, depletion of Nrg led to a significant suppression of the age-related increase in ISC proliferation when compared to outcrossed controls (Supplementary Figure 3A-B). Interestingly, depletion of *Nrg* in the midgut using the RU-inducible *5966^GS^* GeneSwitch ‘driver’, which is expressed primarily in ECs in young flies and in both EB-like cells and ECs in intestines from aged flies (Supp. Fig. 3E-F), also suppressed the increase in ISC proliferation associated with age (Supplementary Figure 3C-D, E-F). Taken together, these data suggest that expression of *Nrg* in EBs in young flies and in EB-like cells in the guts of aged flies could play an important role in altered ISC behavior and loss of intestinal homeostasis over time.

### *Nrg* induces ISC proliferation through EGFR pathway

As noted above, although Nrg is commonly thought to act as a SJ protein, its role in ISCs and EBs is not likely to be in mediating cell-cell adhesion and paracellular flux. Interestingly, previous studies showed that Nrg and hL1CAM can activate signaling via receptor tyrosine kinases such as EGFR and FGFR (Donier et al., 2012; García-Alonso et al., 2000; Islam et al., 2004; Kulahin et al., 2008; Nagaraj et al., 2009). Indeed, genetic analyses have shown that EGFR is acts downstream Nrg in the *Drosophila* brain and that the Nrg-EGFR pathway acts to control growth cone decisions during sensory axon guidance and axonal pathfinding during wing development (García-Alonso et al., 2000; Islam et al., 2004). Furthermore, it has been shown that in S2 cells EGFR and Nrg interact physically, in *trans* and *cis*-configurations, which result in Nrg-mediated activation of EGFR in the absence of classic EGFR ligands (Islam et al., 2004). The interaction between Nrg and EGFR is notable because the EGFR signaling pathway plays a primary role in regulating ISC proliferation and maintenance in the adult fly midgut. EGFR is essential for ISC proliferation under homeostatic conditions, as well as in response to stress signals (Biteau and Jasper, 2011; Buchon et al., 2010; Jiang and Edgar, 2009; Jiang et al., 2011; Jin et al., 2015; Wang et al., 2014a; Xu et al., 2011). Importantly, phenotypes caused by *Nrg* / *L1CAM* over-expression in ISC/EBs (Fig. 3A) are similar to the phenotypes reported for activation of EGFR activity in ISCs: increased proliferation and accumulation of EB-like cells (Biteau and Jasper, 2011; Xu et al., 2011) (Fig. 3B).

To determine whether Nrg activates EGFR signaling in ISC/EBs, we monitored the activity of the EGFR signaling pathway by detecting the levels of the active diphosphorylated form of ERK (dpERK) (Biteau and Jasper, 2011; Gabay et al., 1997; Xu et al., 2011) (Fig. 5A-D). We quantified dpERK intensity in *esg*-positive (GFP^+^) cells in guts of flies overexpressing *Nrg* in ISCs/EBs for 7 days and compared the levels of activation to expression of a *w^1118^* outcross (Fig 5.A-D). Expression of an activated form of EGFR, *EGFR^λtop^*, which signals independently of any ligands (Fig 5. B-B’, D), provided a positive control for dpERK staining. Our analysis showed a similar increase in dpERK intensity when comparing ectopic expression of *Nrg* and *EGFR^λtop^* to controls (Fig. 5A-D).

**Figure 5.**
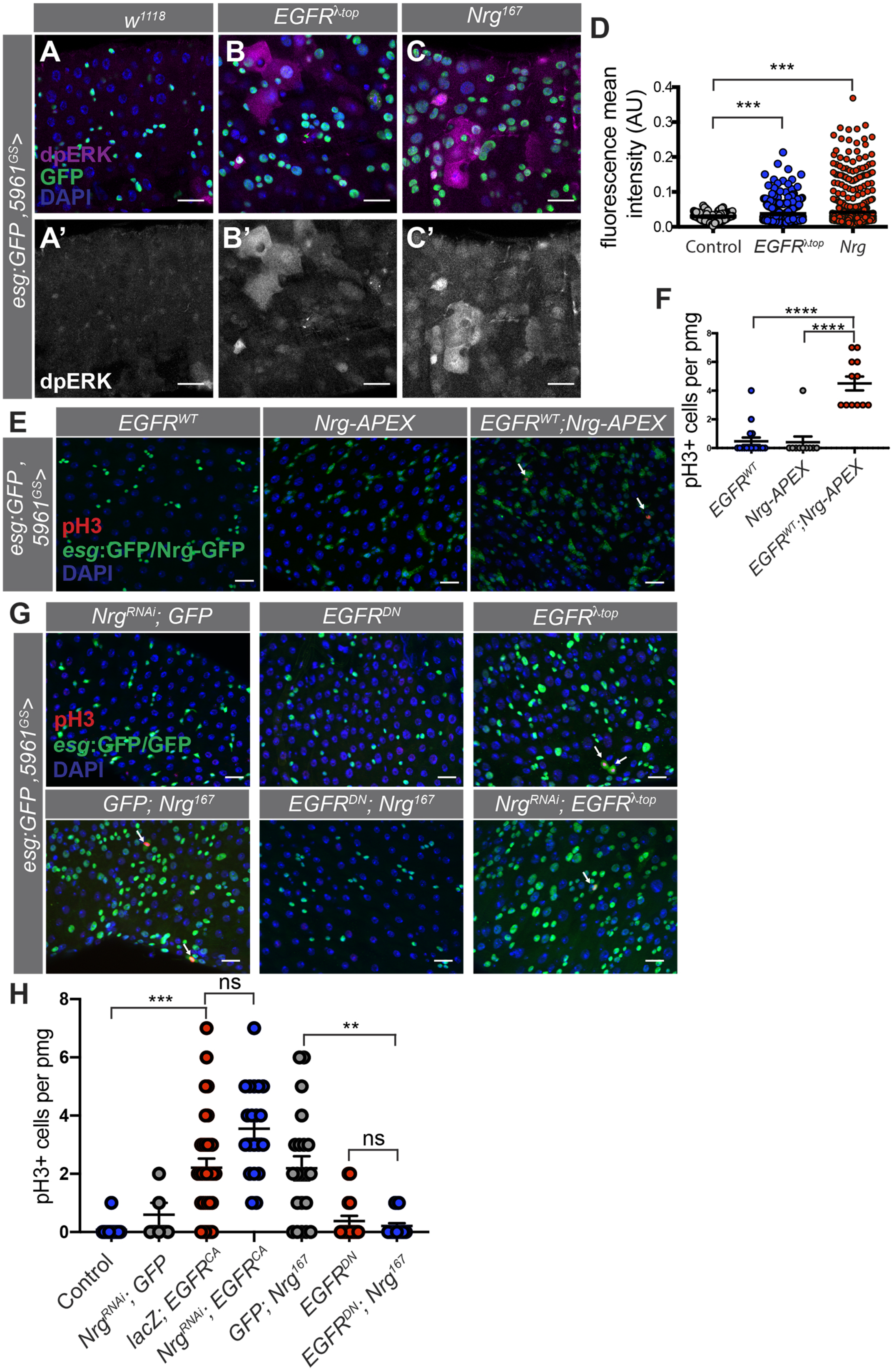
Nrg interacts with EGFR to potentiate signaling. A-C) Images of the target of EGFR signaling dpERK (magenta/grey) in midguts expressing *w^1118^* (A, A’)*, EGFR^λtop^*(B, B’)*, Nrg^167^* (C, C’) driven by *5961^GS^* for 7d. D) Quantification of nuclear dpERK intensity in GFP+ DAPI+ positive cells in A-C. N = 108 *w^1118^* cells, 868 *EGFR^λtop^* cells, 561 *Nrg^167^* cells. E) Representative images from adult midguts of *EGFR^WT^, Nrg^167^-APEX-GFP*, and *EGFR^WT^; Nrg^167^-APEX-GFP* together driven by *5961^GS^* for 7 days. ISC/EBs *(esg:GFP*,green), mitotic cells (pH3, red), and nuclei (DAPI, blue) are shown. F) Quantification of number of mitotic cells per posterior midgut marked by pH3 in A per two fields of view in the pmg in E. N = 17 *EGFR^WT^*, 10 *Nrg^167^-APEX-GFP*, 12 *EGFR^WT^; Nrg^167^-APEX-GFP*midguts. Kruskal-Wallis test with multiple comparisons. G) Representative images from adult midguts from epistasis analysis of *Nrg* and *EGFR* driven by *5961^GS^* for 7 days. ISC/EBs *(esg:GFP*, green), mitotic cells (pH3, red), and nuclei (DAPI, blue) are shown. H) Quantification of number of mitotic cells per posterior midgut marked by pH3 in G per two fields of view in the pmg. N = 13 control, 5 *Nrg^RNAi^; GFP*, 33 *lacZ; EGFR^λtop^*, 20 *Nrg^RNAi^; EGFR^λtop^*, 21 *GFP; Nrg^167^*, 16 *EGFR^DN^*, 19 *EGFR^DN^; Nrg^167^* midguts. Kruskal-Wallis test followed by multiple comparisons.

Unlike EGFR^λtop^, wild type EGFR acts in a ligand-dependent manner (Guichard et al., 1999). To test whether Nrg could enhance activation of wild type EGFR, we coexpressed a GFP-tagged form of Nrg *(Nrg-APEX-GFP)* with wild-type EGFR *(EGFR^WT^)* with *5961^GS^* to express in ISCs and EB. No increase in pH3 was observed as a consequence of *EGFR^WT^* expression or expression of *Nrg-APEX-GFP* (Fig. 5E-F). However, co-expression of *Nrg-APEX-GFP* together with *EGFR^WT^* led to a significant increase in ISC proliferation after 7 days of induction (Fig. 5E-F), indicating that Nrg can potentiate EGFR activation to drive ISC proliferation.

Next, we wanted to determine whether Nrg acts up or downstream of EGFR to stimulate ISC proliferation. As previously observed, overexpression of *Nrg^167^* or activated *EGFR^λtop^* was sufficient to induce ISC proliferation (Figs 3C-D, 5G) (Biteau and Jasper, 2011; Buchon et al., 2010; Jiang et al., 2011; Wang et al., 2014a; Xu et al., 2011) Also as expected, suppression of EGFR signaling in ISC/EBs for 7 days, achieved by ectopic expression of a dominant-negative version of EGFR, *EgfrD^N^*, had no observable effect on ISC/EB proliferation in intestines from young flies due to predictably low levels of proliferation (Fig. 5G) (Biteau and Jasper, 2011; Xu et al., 2011). However, expression of *EgfrD^N^* was sufficient to suppress the increase in ISC proliferation in response to ISC/EB-specific overexpression of *Nrg^167^* (Fig. 4G). In contrast, RNAi-mediated depletion of *Nrg* did not suppress the increase in ISC division caused by *EGFR^λtop^.* Altogether, these data indicate that EGFR signaling is activated downstream of Nrg. In addition, our data suggest that Nrg potentiation of EGFR signaling is important for the proper regulation of ISC behavior in young flies. Thus, we conclude that the increase in Nrg-expressing EB-like cells in intestines of aged flies likely contribute to an increase in ISC proliferation by enhancing EGFR activation, which contributes to the loss of gut homeostasis over time.

## Discussion

Nrg has been characterized previously for its signaling and cell adhesion roles in neural development (Enneking et al., 2013; Goossens et al., 2011; Kristiansen et al., 2005; Kudumala et al., 2013; Moscoso and Sanes, 1995). Additional work, including studies from our lab, have described expression and possible roles for Nrg in SJs in the hindgut and other epithelial tissues (Baumann, 2001; Bergstralh et al., 2015; Genova and Fehon, 2003; Resnik-Docampo et al., 2017; Wei et al., 2004). Here, we show that Nrg is a marker of ISCs and EBs in *Drosophila* and that Nrg plays a significant role in maintaining intestinal homeostasis. Although other SJ proteins that are expressed in ECs have been shown to regulate ISC behavior in a non-autonomous manner (Chen et al., 2020; Resnik-Docampo et al., 2017; Salazar et al., 2018; Xu et al., 2019), the restriction of Nrg expression to ISC/EB nests (Figure 1 and S1) (Baumann, 2001), together with the absence of SJs between ISCs and other progenitor cells, indicated another role for Nrg in the *Drosophila* midgut.

Using clonal analysis and RNAi-mediated depletion we have identified Nrg as a novel regulator of ISC proliferation (Fig. 2, Supplementary Figure 2). Furthermore, ectopic expression of either Nrg or its human homolog, hL1CAM, was sufficient to induce proliferation and accumulation of cells expressing the ISC/EB marker, Esg (Figure 3). Interestingly, ectopic expression in EBs alone using *Su(H)^ts^* was sufficient to cause ISC proliferation while expression in ISCs alone was not (Figure 3), indicating Nrg can act in a non-autonomous manner to stimulate ISC proliferation.

Consistent with expression in EBs and the expansion of EB-like cells with age, an increase in Nrg expression was observed in intestines from aged flies (Figure 4, Supplementary Figure 3). Supporting the idea that the increase in Nrg-expressing cells can drive age-related ISC proliferation, depletion of *Nrg* from EB-like cells was also sufficient to suppress an increase in ISC proliferation in aged flies (Figure 4, Supplementary Figure 4).

In both neurons and epithelial cells in flies, Nrg has been shown to interact with and potentiate the signaling of receptor tyrosine kinases such as the EGFR and FGFR (García-Alonso et al., 2000; Islam et al., 2004). Numerous studies have indicated that EGFR signaling is an essential regulator of ISC proliferation during homeostasis, stress conditions, and aging (Biteau and Jasper, 2011; Buchon et al., 2010; Jiang et al., 2011; Jin et al., 2015; Xu et al., 2011; Zhang et al., 2019); therefore, we tested whether Nrg was important for EGFR signaling in the intestine. Consistent with its role in other tissues, we found that Nrg in ISCs/EBs of the *Drosophila* midgut acts together with EGFR to regulate mitogenic signaling (Figure 5). Therefore, our data support a model in which increases in EB-like cells with age would lead to increased Nrg, which in turn exacerbates EGFR signaling, resulting in uncontrolled ISC divisions and, ultimately, intestinal dysplasia.

Previous research has shown that Nrg/hL1CAM can signal through both heterotypic and homotypic interactions at cell-cell contacts to potentiate signaling (Donier et al., 2012; Enneking et al., 2013; Islam et al., 2004). Further, Nrg was capable of activating EGFR/Erk signaling in the absence of additional ligands *in vitro* (Islam et al., 2004). In the *Drosophila* midgut, the EGFR ligands Vein (secreted by visceral muscle), and Keren/Spitz (from progenitors and ECs), have been described to stimulate ISC proliferation primarily during homeostasis and stress (Biteau and Jasper, 2011; Buchon et al., 2010; Jiang et al., 2011; Patel et al., 2015; Xu et al., 2011). Limited data is available on the participation of the various ligands in EGFR activation with aging. Therefore, further research is needed to determine whether Nrg may activate EGFR independently, or in conjunction with, traditional agonists.

Intriguingly, recent research has identified age-related disruption of the endocytosis/autophagy pathway as one mechanism leading to an increase in EGFR and, consequently Erk signaling in aged ISCs via stabilization of ligand-activated EGFR (Du et al., 2020; Zhang et al., 2019). Our work suggests that an increase in Nrg-mediated potentiation of EGFR signaling may be additional mechanism that contributes to increases in ISC proliferation and intestinal dysplasia with age (Figure 4, Supplementary Figure 4).

Increased hL1CAM expression is associated with a variety of cancers (Altevogt et al., 2016; Gavert et al., 2008) and tumor metastasis (Ernst et al., 2018; Fang et al., 2020; Gavert et al., 2010; Huszar et al., 2010; Lund et al., 2015; Terraneo et al., 2020), including gastrointestinal cancers (Fang et al., 2020; Ganesh et al., 2020; Gavert et al., 2005, 2010). Mechanistic studies have shown that increases in hL1CAM may be associated with EMT to drive metastasis (Ernst et al., 2018; Giordano and Cavallaro, 2020; Huszar et al., 2010; Lund et al., 2015; Tischler et al., 2011; Versluis et al., 2018). Additionally hL1CAM was required for growth and proliferation of intestinal organoids derived from colorectal cancer (CRC) tissue (Ganesh et al., 2020). Indeed, increases in hL1CAM have been shown to regulate CRC metastasis via ERK signaling (Fang et al., 2020), indicating that the relationship between Nrg-EGFR may be conserved in the mammalian intestine.

Therefore, a better understanding of the role of Nrg/hL1CAM-EGFR signaling in stem cell proliferation and maintenance may lead to the development of additional strategies to target this pathway in the initiation and progression of CRC and disorders caused by excess EGFR activation.

## Acknowledgements

The authors thank H. Jasper (Genentech and The Buck Institute for Research on Aging, USA), Ben Ohlstein (Columbia University, USA), J. Pielage (University of Kaiserslautern, Germany), J. F. de Celis (CBMSO, Spain), A. Baonza (CBMSO, Spain), G. Tanentzapf, (University of British Columbia, Canada), the Vienna Drosophila RNAi Center (VDRC), Kyoto Stock Center (DGRC) and Bloomington Stock Center for reagents, the BSCRC/MCDB microscopy core at UCLA, and the Jones laboratory for comments on the manuscript.

## Experimental Procedures

### Fly food and husbandry

All analyses for these studies were performed on female flies, as age-related gut pathology has been well established in females (Biteau et al., 2008; Rera et al., 2012). Flies were cultured in vials containing standard cornmeal medium (1% agar, 3% brewer’s yeast, 1.9% sucrose, 7.7% molasses or 7.8% malt syrup, and 9.1% cornmeal; all concentrations given in wt/vol). The drug-inducible GAL4 ‘Geneswitch’ (Osterwalder et al., 2001; Roman et al., 2001) system was used with the drivers *Su(H)lacZ; esg:GFP,5961-GAL4^GS^* or *Su(H)lacZ; esg:GFP, 5961-GAL4^GS^* crossed to *UAS-Nrg^RNAi^* or *UAS-Nrg^167^* or outcrossed to control *(UAS-mCherry^RNAi^* or *w^1118^*) and raised at 25 °C on standard food. Progeny were collected at eclosion and allowed to mate and develop for 3-5 days before being transferred to food mixed with 50ug/mL mifepristone (RU486, Sigma) or ethanol (control) at 25 °C for induction time noted. Flies were transferred to new food vials every 2-3 days thereafter. Aged flies in Figure 4D-E were induced for the entire aging interval. For temperature sensitive (ts) crosses using GAL80^ts^, crosses were set and maintained at 18°C until eclosion. Afterwards, adults were kept for 2-3 days at 18°C and then moved to 29°C for induction as noted.

### Fly lines

Lines not described in the text can be found in Flybase.

| Stock | Source | Identifier |
| --- | --- | --- |
| *Su(H)lacZ; esg:GFP,5961-GAL4<sup>GS</sup>* | Gift from B. Ohlstein, Columbia University, USA (Mathur et al., 2010) |  |
| *Su(H)lacZ; esg:GFP,5966-GAL4<sup>GS</sup>* | Gift from B. Ohlstein, Columbia University, USA |  |
| *Su(H)-GAL4, UAS-CD8GFP; tubGAL80<sup>ts</sup>* | Gift from S. Hou (Zeng et al., 2010) |  |
| *esg-GAL4, UAS-2xYFP; Su(H)GAL80, tubGAL80<sup>ts</sup>* | Gift from S. Hou (Wang et al., 2014b) |  |
| *SalEPGAL4* | Gift from J. F. de Celis |  |
| *y,w,hsFLP1.22 P[tub-gal4] UAS-GFP; P[tub-gal80<sup>ts</sup>] FRT40A/CyO* | Gift from A. Baonza, CBMSO, Spain |  |
| *Nrg*<sup>HMS01638</sup> Valium 20 |  | BDSC 37496 |
| *Nrg*<sup>GD14467</sup> |  | VDRC 27201 |
| *Nrg*<sup>14</sup> FRT19A | Gift from Jan Pielage<br>Division Zoology-Neurobiology<br>Technische Universität<br>Kaiserslautern |  |
| *Nrg*<sup>G00413</sup> FRT19A |  | Kyoto stock center 111923 |
| *UAS-Nrg*<sup>167</sup> |  | BDSC 24172 |
| *UAS-hL1CAM* | (Islam et al., 2004) | BDSC 24171 |
| *UAS-EGFR*<sup>λ<sub>top</sub></sup> |  | BDSC 59843 |
| *UAS-EGFR*<sup>DN</sup> |  | BDSC 5364 |
| *UAS-EGFR*<sup>WT</sup> |  | BDSC 5368 |
| *Neuroglian::GFP*<br>(*Nrg::GFP*) | Gift from G. Tanentzapf<br>University of British<br>Columbia, Canada |  |

### Generation of Neuroglian antibody

The Neuroglian antibody was designed and generated by Thermofischer Scientific. A synthetic peptide from the C-terminal *Nrg* sequence 1204:1222: KPGVESDTDSMAEYGDGDT was generated and used to inject rabbits. Anti-sera were collected after 96 days. Unpurified sera were used for staining.

### Fluorescence microscopy and antibody staining

For consistency, imaging was always done on the P3–P4 regions of the *Drosophila* intestine, located by centering the pyloric ring in a ×40 field of view (fov) and moving 1– 2 fov toward the anterior. Posterior midguts were dissected into ice-cold phosphate buffered saline (PBS)/4% paraformaldehyde (PFA) and incubated for 1 h in fixative at room temperature. Samples were then washed three times, for 10 min each, in PBT (PBS containing 0.1% Triton X-100), and incubated in block (PBT-0.5% or PBT-0.3% bovine serum albumin) for 30 min. Samples were immunostained with primary antibodies overnight at 4 °C, washed 4× 5 min at room temperature in PBT, incubated with secondary antibodies at room temperature for 2 h, washed three times with PBT and mounted in Vecta-Shield with DAPI (Vector Laboratories, H-1200).

For anti-dp-ERK stainings, the following protocol was modified from (Castanieto et al., 2014). Flies were placed on food supplemented with yeast paste overnight prior to dissection. Posterior midguts were dissected into ice-cold PBS with phosphatase inhibitor (1:100, Sigma, cat#P5726). Guts were in ice cold PBS/4% PFA with phosphatase inhibitor and then taken through a methanol (MeOH) dehydration as follows: 25% MeOH 3 min, 50% MeOH 3 min, 75% MeOH 3 min, 100% MeOH 3 min, 75% MeOH 3 min, 50% MeOH 3 min, 25% MeOH 3 min. All MeOH solutions contained phosphatase inhibitors Guts were then washed three times, for 10 min each, in PBT (PBS containing 0.3% Triton X-100) plus phosphatase inhibitor. Guts were blocked and immunostained as above with the addition of phosphastase inhibitor in all solutions.

Primary antibodies used include: rabbit anti-GFP (1:3,000, Molecular Probes A-11122); mouse anti-GFP (1:200, Molecular Probes A-11120); chicken anti-GFP (1:500, Aves Labs GFP-1010); rabbit anti-β-GAL (1:2,000, Cappel/MPbio 559761); rabbit anti-Phospho-histone3 (1:200, Millipore 06-570). Rabbit anti-dsRed (1:100, Clontech, 632496), Rabbit anti-Neuroglian (1:50, this study); mouse anti-EGFR (1:1000, Millipore Sigma E2906); rabbit anti-phospho-p44/42 MAPK (1:100 Cell Signaling Technologies Cat #4370). The Armadillo antibody used (mouse, 1:100) was obtained from the Developmental Studies Hybridoma Bank, developed under the auspices of the NICHD and maintained by The University of Iowa, Department of Biology, Iowa City, IA 52242:.

Images were acquired on a Zeiss LSM710 or LSM800 inverted confocal microscope, and/or on a Zeiss Axio Observer Z1, and processed with Fiji/ImageJ (NIH) and Zen Blue or Black software (Zeiss). The final figures were assembled using Adobe Photoshop or Adobe Illustrator.

### Generation of MARCM clones

Mutant clones: *hs-flp,tubGal80, neoFRT19A; UAS-mCD8::GFP* flies were crossed to *FRT19A, Nrg^14^/FM7; P[tub-Gal4]/CyO* or *FRT19A, Nrg^G00413^/FM7; P[tub-Gal4]/CyO* or *FRT19A/FM7; P[tub-Gal4]/CyO* (control) flies, and progeny raised at 25°C were treated heat shocked at 37°C for 90 minutes once or twice on the same day, 6-7 hours apart. The flies were then placed back at 25°C and dissected at designated time points, as noted in figure legends (Fig 2).

RNAi clones: *y,w,hsFLP1.22 P[tub-Gal4] UAS-GFP; P[tub-Gal80^ts^] FRT40A/CyO* flies were crossed to *FRT40A/CyO; UAS-Nrg^RNAi GD^/TM6B* or *FRT40A/CyO; 2xUAS-GFP/TM6B* (control). In order to generate clones, progeny raised at 25°C were heat shocked at 37°C for 90 minutes twice on the same day, 6-7 hours apart. The flies were then placed back at 25°C and dissected at designated time points, as noted in figure legends (Fig S2).

### Generation of *UAS-Nrg-APEX2-GFP*

APEX2-EGFP (gift from M. Ellisman) was inserted into the vector ‘pUASt attBK7 SfiI BglII EcoRI’ (a gift from M. Rera and D. Walker, UCLA). Full-length Nrg cDNA obtained from DGRC (clone GH03573) was inserted into the linearized backbone (EcoRI, SfiI) using an In-Fusion HD Cloning Kit (Takara Bio). Site-specific attp40 insertion into the fly genome was performed by Bestgene, Inc.

### Quantification of Nrg levels in intestines from aged flies

Posterior midguts from 3 or 45 day old *Nrg::GFP* flies were analyzed. ISC/EB ‘nests’ in the posterior region of the midgut were imaged, as described above. All possible “nests” were imaged in the field of view, and the total number of Nrg::GFP^+^ cells and DAPI+ cells per field of view were counted by eye in ImageJ. 33 fields of view from 8 young intestines and 41 fields of view from 9 old intestines were analyzed.

### Quantification using CellProfiler.

Analysis was conducted as described in (Resnik-Docampo et al., 2017). Briefly, two z-stacks with a typical slice thickness of 750 nm were taken from the same side of each posterior midgut (using a minimum of 6 guts). The images were then processed using CellProfiler (Carpenter et al., 2006; Kamentsky et al., 2011) to automatically quantify dpERK intensity cells within GFP^+^ cells. Average ratios from the two images corresponding to a single gut were used in subsequent statistical analyses.

### Statistics and reproducibility

Statistical analysis and graphical display of the data were performed using Prism6 (GraphPad). Significance, expressed as P values, was determined with a two-tailed test; the following parametric or non-parametric tests were used as indicated in the figure legends: One-way ANOVA/Tukey’s multiple comparisons test or Student’s t-test were used when data met criteria for parametric analysis (normal distribution, equal variances), Kruskal-Wallis/Dunn multiple comparisons test was used when data were non-parametric. Experiments were repeated at least two times. No statistical method was used to predetermine sample size. The experiments were not randomized and investigators were not blinded to allocation during experiments and outcome assessment.

### Data availability

RNA sequencing data were previously published (Resnik-Docampo et al., 2017) and deposited in the Gene Expression Omnibus (GEO) under the accession number <u>GSE74171</u>. All other data supporting the findings of this study are available from the corresponding author on request.

