## Supplemental Figures and Figure Legends for "Neuroglian regulates *Drosophila* intestinal stem cell proliferation through enhanced signaling via the Epidermal Growth Factor Receptor"

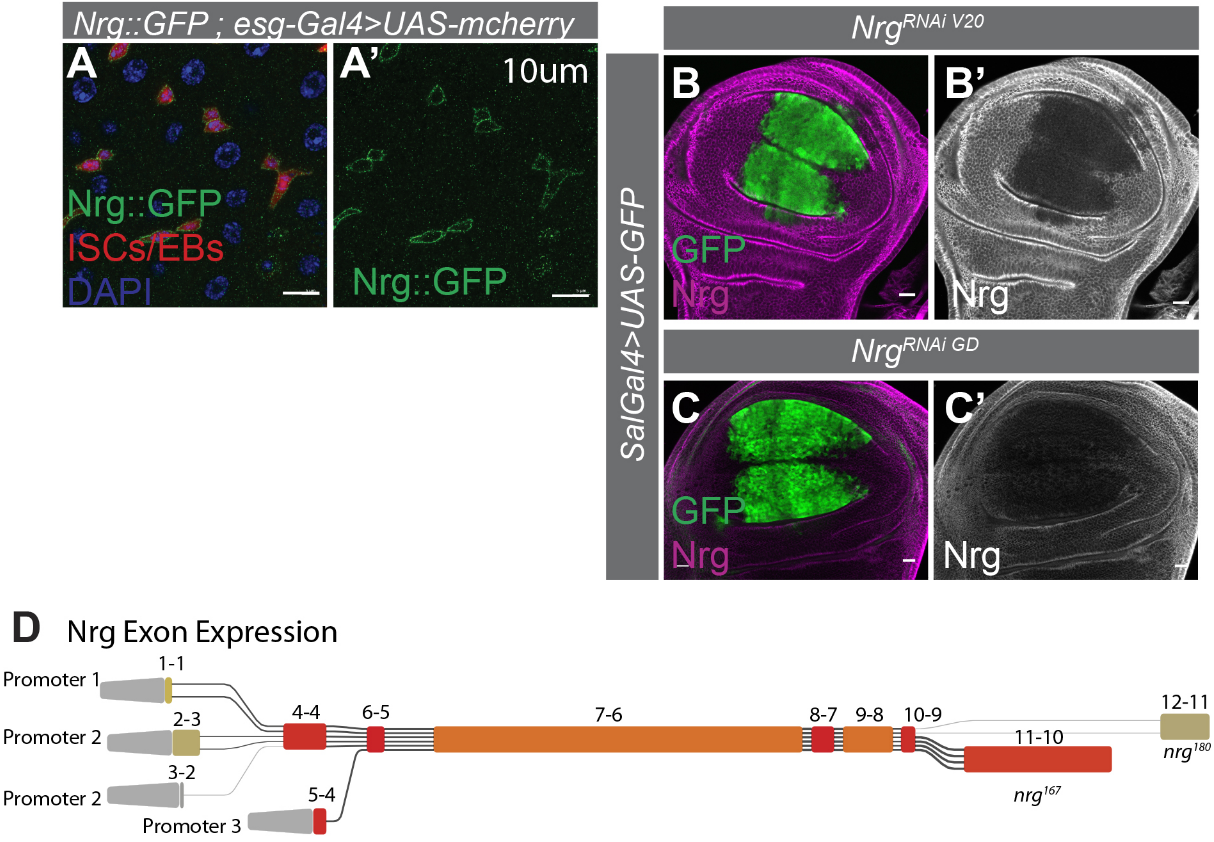


**Supplemental Figure 1**

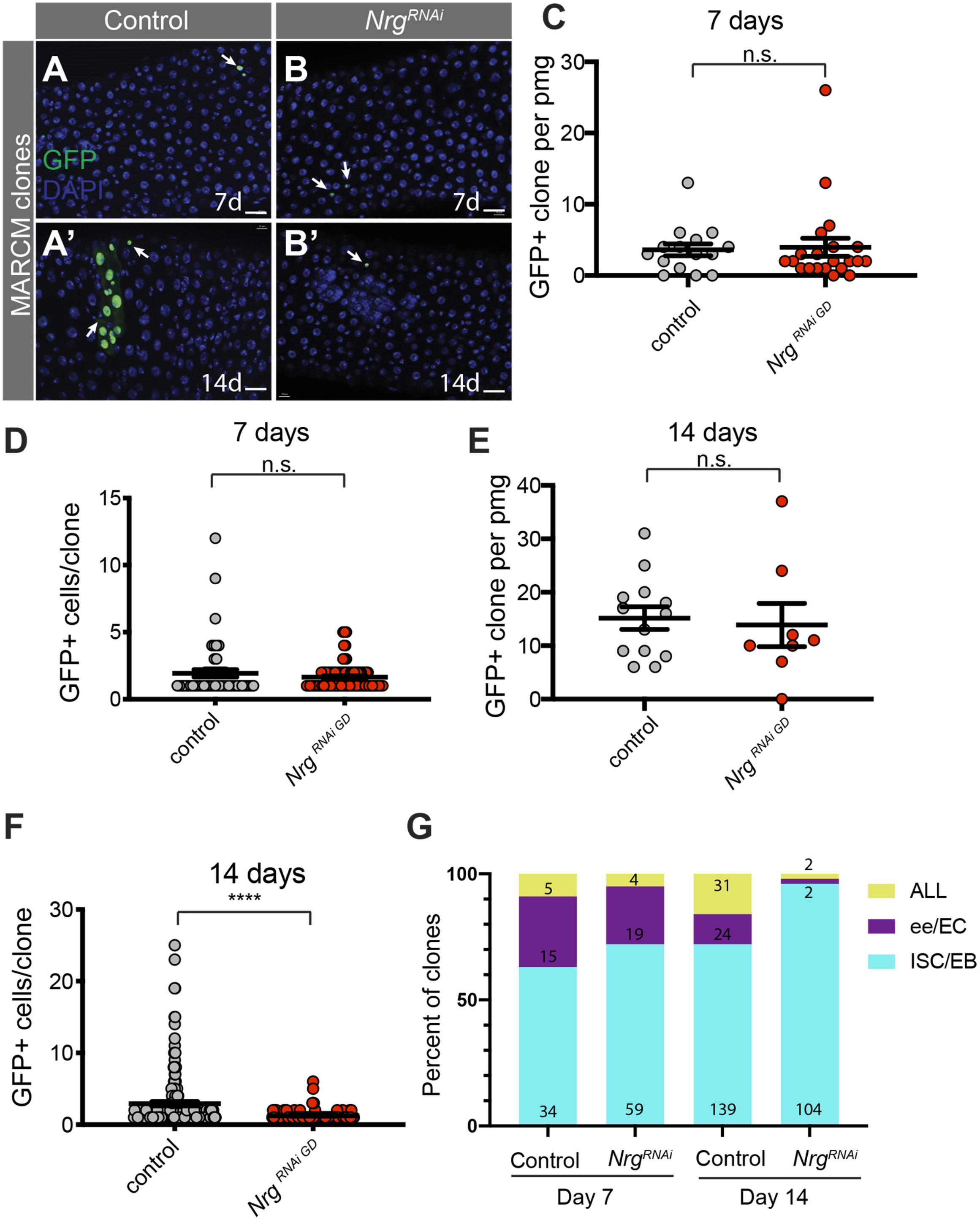


**Supplemental Figure 2**

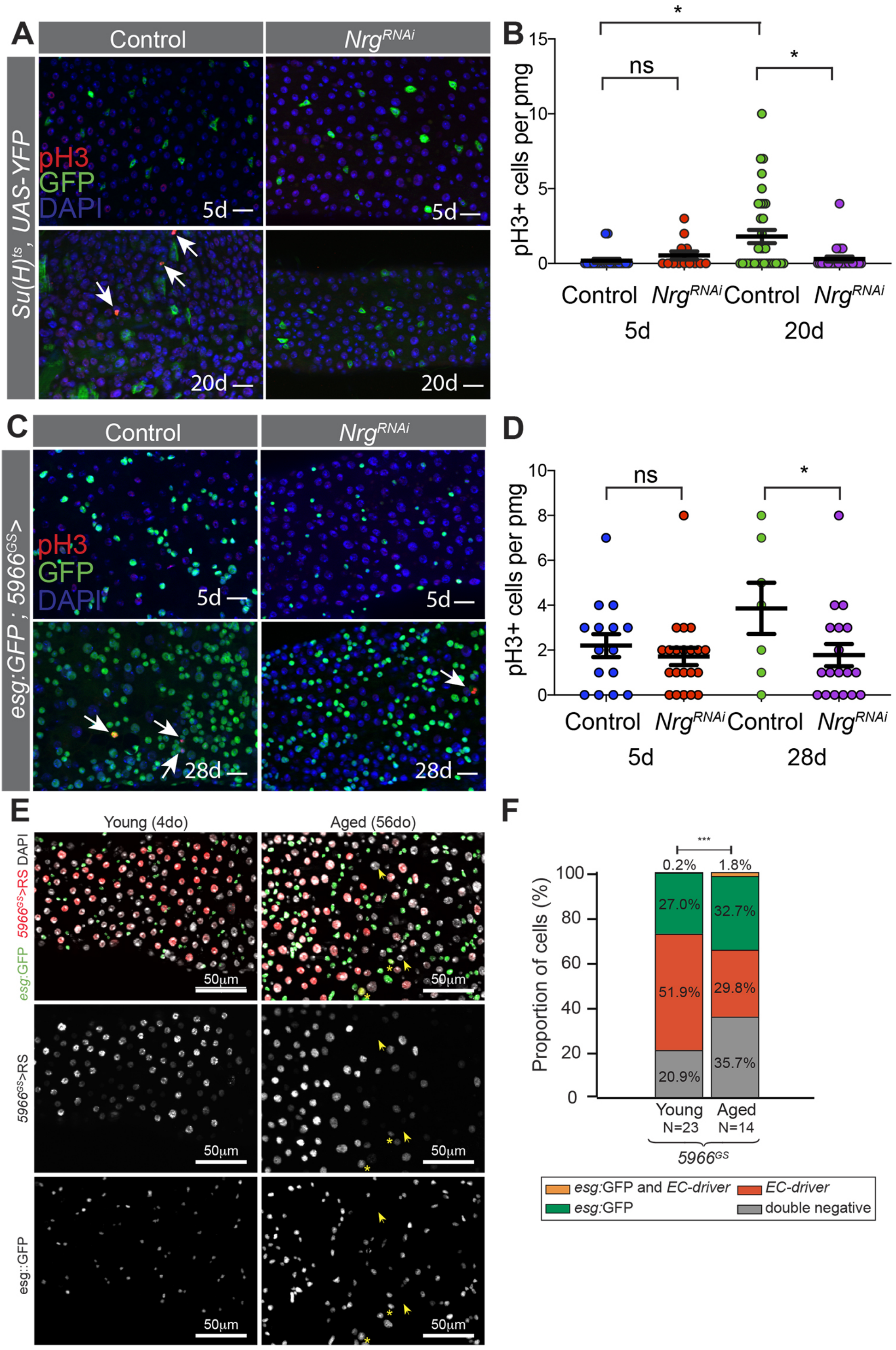


**Supplemental Figure 3**

**Supplementary Figure Legends**

**Figure S1 (Related to Figure 1). Nrg is expressed in the ISCs/EBs of the *Drosophila* midgut.** A) Image of *Drosophila* midguts with *Nrg::GFP* (A') and *esgGal4>mcherry*. Note co-localization in ISC/EB nests. Scale bar = 10μm) B-D) *SalGal4>GFP* in the wing disc driving $Nrg^{RNAi}$ V20 (B-B') or $Nrg^{RNAi}$ GD (C-C') stained with anti-Nrg antibody. *Nrg* is depleted specifically in GFP+ cells expressing the RNAi. Scale bar = 20μm. D) Heat map of exon expression levels of *Nrg* in young midguts detected by RNA-Seq (see methods). Note high expression of exon corresponding to the $Nrg^{167}$ isoform.

**Figure S2 (Related to Figure 2). Nrg is required for ISC proliferation.** Examples of midguts 7 and 14 days after MARCM clonal generation in control (A, A') and $Nrg^{RNAi}$ (B, B') backgrounds. Scale bar = 20μm. C) Quantification of GFP+ clones per midgut after 7 days in A-B. N = 15 (control guts), 21 ($Nrg^{RNAi}$ guts), Mann-Whitney test D) Quantification of GFP+ cells per clone after 7 days in A-B. N = 54 (control clones), 81 (clones expressing $Nrg^{RNAi}$), Mann-Whitney test. E) Quantification of the number of GFP+ clones per midgut after 14 days in A'-B' N= 13 (control guts), 8 ($Nrg^{RNAi}$ guts), Mann-Whitney test F) Quantification of GFP+ cells per clone after 14 days in A'-B' N = 197 (control clones), 109, (clones expressing $Nrg^{RNAi}$), Mann Whitney test. G) Characterization of the types of cells per clone (ISC/EB progenitor, ee/EC differentiated cells, or ALL progenitors and differentiated cells) marked by GFP in A-B.

**Figure S3 (Related to Figure 4)** A) Representative images of midguts expressing *Su(H)Gal4, UAS-GFP; tubal80*$^{ts}$*> Nrgi V20* or outcross control ($w^{1118}$) either 5 or 20d after induction at 29C, stained for GFP (*Su(H)>GFP*, green), pH3 (red), and DAPI. B)

Quantification of the number of pH3+ cells per pmg in A, N= 22 (control 5do), 13 (*Nrg*$^{RNAi}$ 5do), 35 (control 20do), 27 (control 20do) midguts, Kruskal-Wallis test with multiple comparisons. Note: Controls for day 5 the same as for the Panel 3F-G, and experiments were conducted in parallel. C) Representative images of midguts expressing *5966*$^{GS}$> *Nrg RNAi GD* or outcross control (*w*$^{1118}$) after 5 or 28d feeding with 50ug/mL RU486. D) Quantification of the number of pH3+ cells per pmg in C, N= 15 (control 5do), 21 (*Nrg*$^{RNAi}$ 5do), 7 (control 28do), 18 (*Nrg*$^{RNAi}$ 28do). E) Expression pattern of the driver *5966*$^{GS}$*>UAS-RedStinger* in midguts of young (4 d.o.) and old (56 d.o.) flies. Asterisks indicate cells where the ISC marker esg:GFP and the EC marker *5966*$^{GS}$*>UAS-RedStinger* are both expressed. Arrows indicate DAPI+ cells that are negative for both markers. F) Quantification of the proportion of cells in E single positive for the markers *esg*:GFP or *5966*$^{GS}$*>UAS-RedStinger*, double positive, or double negative (unpaired Student's t-test, N = 23 young midguts, 14 aged midguts).
